# Lipid Defect Analyzer (LipiDeA): A Python Framework for Quantifying Lipid Packing Defects in Curved and Neutral Lipid-Containing Membranes

**DOI:** 10.64898/2026.09.30.755456

**Authors:** Arvin Nazari, Yu-ming M. Huang

**Affiliations:** Department of Physics and Astronomy, Wayne State University, Detroit, MI 48201, USA; Barber Center for Multiscale Systems Biology, Detroit, MI 48202, USA

**Keywords:** Membrane biophysics, Membrane analysis, Lipid organization, Surface reconstruction, Hydrophobic exposure

## Abstract

Lipid packing defects expose hydrophobic regions of membranes and influence protein-membrane recognition, membrane remodeling, and lipid sensing. Quantitative analysis of these defects remains challenging for heterogeneous membranes with strong curvature, mixed bilayer and monolayer organization, and substantial neutral-lipid content. Here, we present Lipid Defect Analyzer (LipiDeA), an open-source Python framework for quantitative analysis of lipid packing defects in complex membrane systems. LipiDeA reconstructs membrane surfaces using triangulated meshes that adapt to local membrane geometry and identifies packing defects from lipid coverage of the reconstructed surface. The framework further classifies bilayer and monolayer regions, resolves defect properties according to local membrane curvature, and distinguishes defects associated with phospholipid tails and neutral lipids. Validation using planar and buckled DPPC bilayers reproduced previously reported packing defect constants and showed that defects preferentially occur in positively curved membrane regions. Application to lipid-droplet-like membranes revealed composition- and topology-dependent defect formation, with the largest defects occurring during the transition from bilayer-dominated to monolayer-containing membranes and distinct defect properties for DAG- and TAG-containing systems. These results highlight the importance of local membrane geometry and composition in determining lipid packing defects.

## 1. Introduction

Biological membranes are essential structural components that define cellular boundaries and provide the physical platform for many biochemical processes. Rather than uniform bilayers, they are dynamic assemblies of diverse lipids whose compositional heterogeneity causes local membrane properties, e.g., thickness, curvature, and lipid order, to fluctuate^1^. At nanoscales, this heterogeneity gives rise to transient voids at the membrane interface where lipid headgroups are imperfectly packed, resulting in exposing portions of the hydrophobic core to the surrounding solvent^2–5^. These interfacial voids, commonly termed lipid packing defects, constitute a fundamental physical property of membrane surfaces. Extensive studies have shown that lipid packing defects regulate diverse membrane-associated processes by creating transient hydrophobic insertion sites at the membrane interface^3,6–12^. Peripheral proteins exploit these defects for membrane recognition, as demonstrated for amphipathic lipid packing sensor motif^13^. Packing defects also facilitate insertion of amphipathic helices, thereby modulating membrane affinity and binding selectivity^14^. Beyond protein recruitment, defect-rich membranes promote membrane remodeling, including tubulation and budding, by lowering the energetic barrier for protein-induced deformation^15–17^. For example, α-Synuclein senses lipid packing defects to initiate helix insertion and drive lateral lipid expansion during membrane remodeling^16^. Lipid packing defects further contribute to lipid sensing and intracellular trafficking by regulating protein localization across membrane compartments^18^.

Several computational approaches have been developed to quantify lipid packing defects from molecular dynamics (MD) simulations. Early analyses treated defects as surface cavities projected onto a 2D Cartesian grid, which enabled systematic characterization of exposed hydrophobic regions in planar lipid bilayers. A notable example is PackMem, which provides an efficient framework to identify shallow and deep defects based on membrane surface topography and has become widely used for analyzing planar bilayer systems^19,20^. Subsequent studies recognized that defect depth is an important descriptor and extended these analyses to 3D representations, which show that defect area alone does not fully capture the structural features relevant for membrane binding^21^. More recently, surface-based methods using triangulated membrane representations have been introduced to evaluate defect distributions in curved membranes and to examine the coupling between local curvature, lipid order, and molecular recognition^22^. The above studies established lipid packing defects as a quantitative membrane descriptor and provided valuable tools for connecting membrane structure to protein binding.

Despite these advances, current methods remain limited by assumptions that are best suited for conventional planar bilayers. Most approaches define packing defects by projecting membrane coordinates onto a Cartesian plane, which implicitly assumes a fixed membrane normal, symmetric leaflets, and interfaces composed primarily of phospholipids^19,20^. These assumptions are reasonable for flat bilayers but become less valid for complex membrane geometries, where local surface orientation varies continuously and the two leaflets may not be equivalent. Although recent triangulated surface methods extend defect analysis to curved membranes, some available implementations are not fully open-source or readily adaptable for custom workflows, which limits broader use and reproducibility^22,23^. These limitations are particularly significant for emerging membrane systems that deviate from the planar bilayer model, including highly curved membranes, membrane tubules, vesicles, and lipid droplets (LD)^3,4,12,24,25^. In addition, neutral-lipid-rich assemblies containing triglycerides (TAG) or diacylglycerols (DAG), as well as monolayer interfaces, present distinct interfacial architectures that challenge bilayer-based definitions of membrane surfaces and defect regions^4,26,27^.

To address these limitations, we developed **Lipi**d **De**fect **A**nalyzer (LipiDeA), a Python program within the **Py**thon **Lip**id and **M**embrane **A**nalysis **P**ackage (PyLipMAP) suite, which provides a collection of analysis tools for lipid and membrane simulations. LipiDeA is a versatile 3D defect analysis tool that identifies hydrophobic packing defects using a triangulated mesh that conforms to the membrane surface. This surface-adaptive strategy enables accurate and efficient characterization of packing defects in both planar and curved membrane systems without requiring a predefined global membrane normal. The current implementation supports coarse-grained (CG) membrane trajectories and is compatible with phospholipid and neutral-lipid systems in both bilayer and monolayer structures. The package is highly customizable and user-friendly. Users can readily adjust a wide range of parameters to suit different membrane compositions, force fields, simulation resolutions, and analysis objectives. LipiDeA also provides visualization outputs that can be directly rendered in VMD and Chimera to facilitate structural interpretation of defect distributions^28,29^. The generated meshes can also be inspected and further processed using dedicated 3D mesh programs, such as MeshLab and Open3D^30,31^. Moreover, the package is implemented using simple and freely available Python libraries, which promotes accessibility and integration into existing analysis workflows. To demonstrate its capability, we apply LipiDeA to both planar and curved membrane systems, including the endoplasmic reticulum with highly curved buckle membranes and LDs containing varying concentrations of neutral lipids. We further compare its results with existing defect analysis tools and assess computational performance to reveal its accuracy, flexibility, and efficiency for diverse membrane geometries.

## 2. Algorithm

LipiDeA is designed to quantify lipid packing defects using a triangulated membrane surface representation. The method operates on 3D coordinates of CG beads and provides a unified framework for defect identification in membrane systems. The workflow consists of 7 independent steps (described below) (**Fig. 1**), each of which can be executed or restarted separately as needed. The entire pipeline is controlled through a single input configuration file. The user can define key analysis parameters, including the force field, bead types used for membrane surface construction, mesh resolution, and clustering size for defect regions. This flexible design enables application to a wide range of membrane compositions and simulation setups.

**Figure 1.**
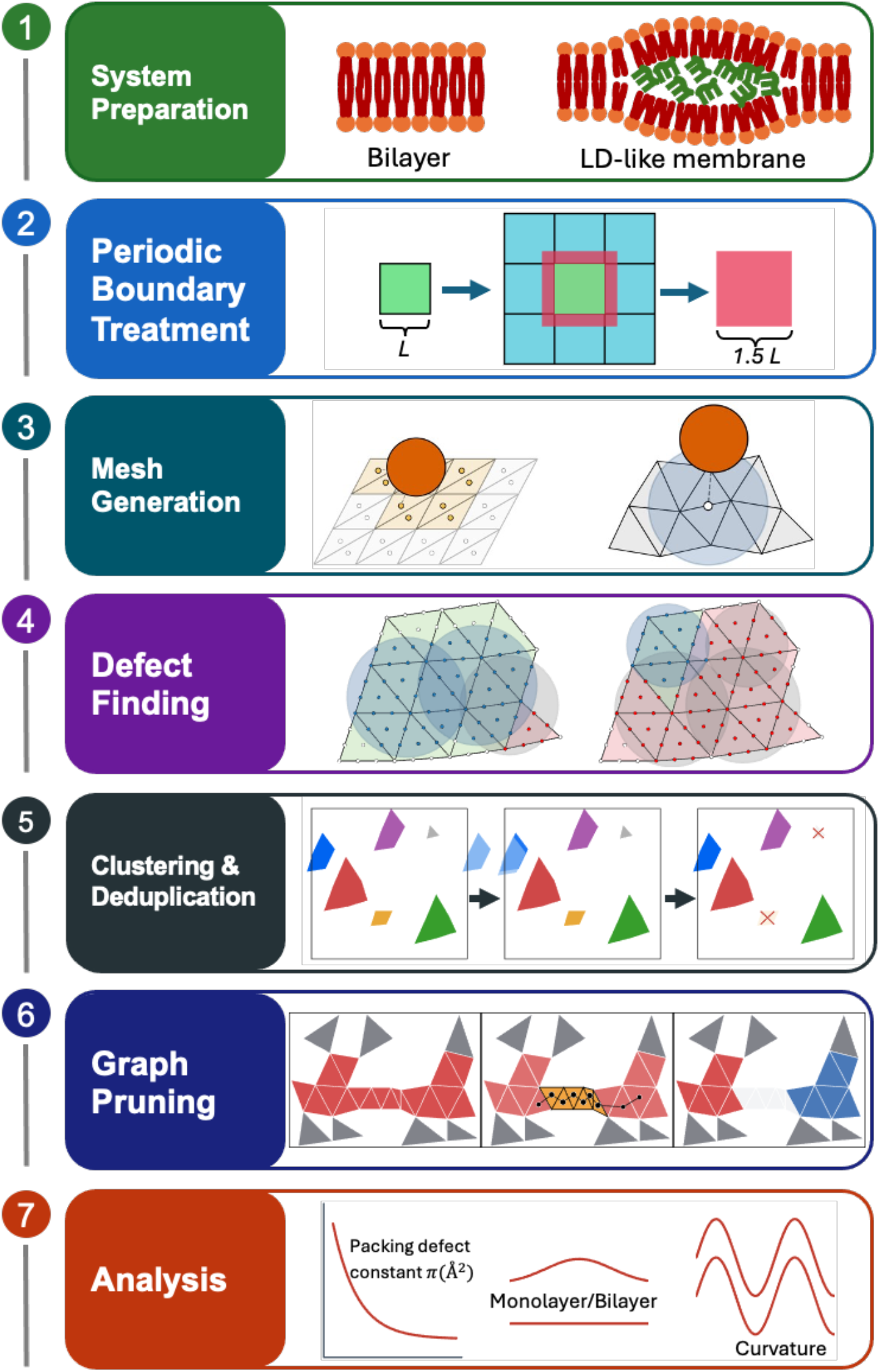
Overview of the LipiDeA workflow. The workflow consists of seven steps. (1) System preparation: bead identities are determined, van der Waals radii are assigned, and the upper and lower membrane leaflets are identified. Phospholipid headgroups, phospholipid tails, and neutral lipids are shown in orange, red, and green, respectively. (2) Periodic boundary treatment: each trajectory frame is replicated on a 3×3 grid in the x-y plane, and a central region with dimensions of 1.5L_x_×1.5L_y_ is extracted. (3) Mesh generation: a triangulated surface is reconstructed independently for each leaflet. (4) Defect finding: van der Waals radii of beads are projected onto the triangle surface. If the headgroup projection covers the triangle, it is considered non-defective; otherwise, it is classified as a defect. Defects from neutral lipids and acyl tails are shown in green and red, respectively. (5) Clustering and deduplication: edge-connected defect triangles are grouped into clusters, shown in different colors. Periodic duplicates are removed, and clusters below the user-defined size threshold are discarded (×). (6) Graph-based pruning: bridge connections (orange) between otherwise distinct defect regions are identified and removed, thereby separating a merged cluster into independent defects. (7) Analysis: the identified defects are characterized through three analyses: packing defect constant calculation, monolayer/bilayer classification, and local membrane curvature analysis.

### 2.1 System Preparation

LipiDeA accepts MD trajectories in PDB format as input. The program processes each trajectory frame independently, which enables efficient analysis of large trajectories. For each frame, bead identities, e.g., van der Waals radii, are assigned according to the selected force field, with MARTINI 3 as the default^32^. The program then classifies membrane constituents into phospholipid headgroups, phospholipid tails, and neutral lipids according to residue and bead identities. LipiDeA records the assigned bead properties and lipid classifications for user inspection. Users may modify these annotations to accommodate custom lipid species or force-field parameters before continuing the analysis.

### 2.2 Periodic Boundary Treatment

To account for periodic boundary conditions, each frame is first replicated in the x-y plane using a 3 × 3 tiling scheme. A central region is then extracted from the replicated system around the original coordinate center. The extracted region spans 1.5 times the original box length in both x and y directions (**Fig. 1(2)**). This process preserves membrane structures that extend across periodic boundaries while reducing edge artifacts in defect identification. At the same time, it avoids the computational cost associated with retaining the full tiled system.

**Figure 2:**
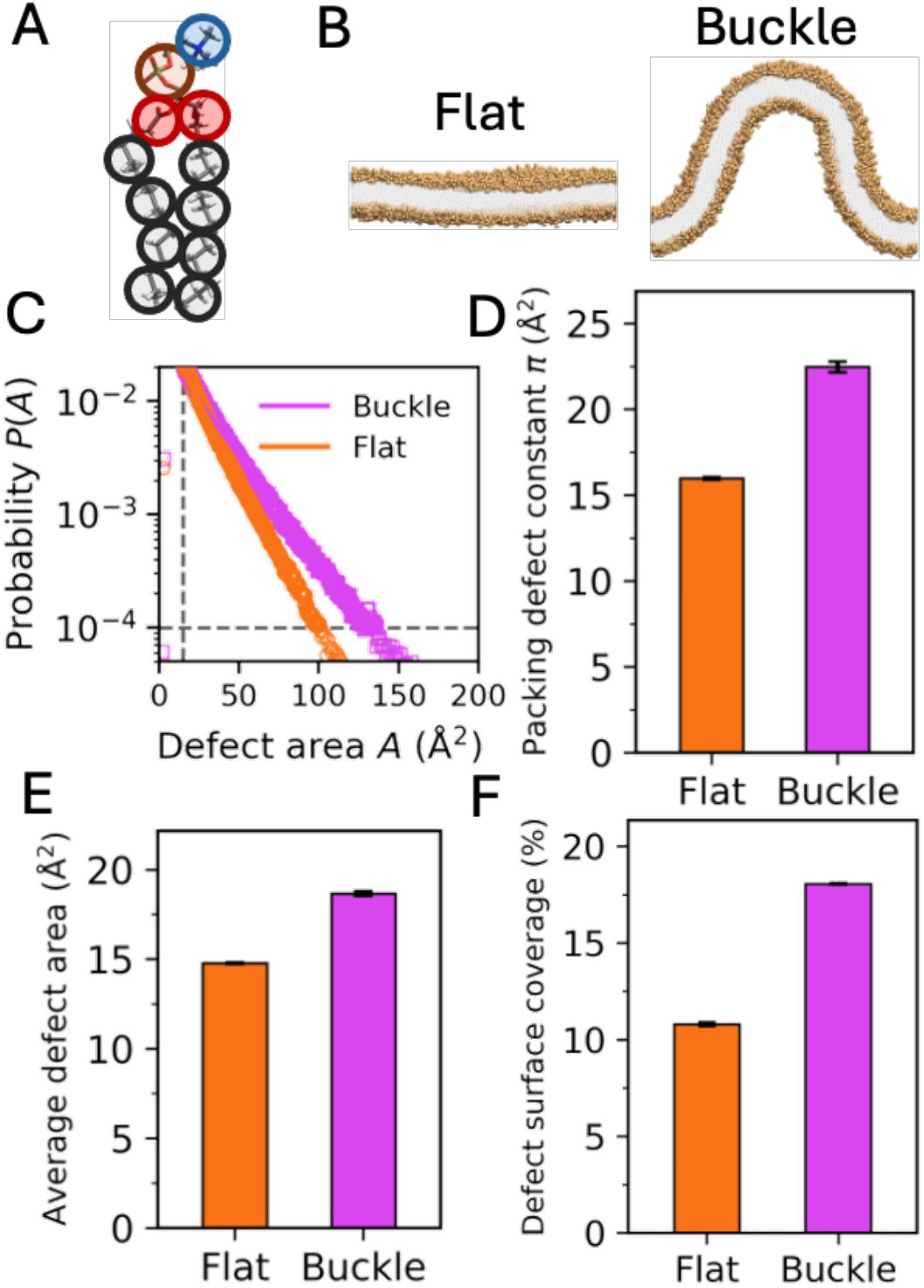
DPPC bilayer systems and their packing defect statistics. (A) Coarse-grained MARTINI bead representation of DPPC compared with all-atom structure. (B) Planar and buckled geometries of the DPPC bilayer. (C) Defect area probability distributions *P(A)* with exponential fits over the 15-250 Å² range and a minimum probability cutoff of 10⁻⁴. (D) Packing defect constant π (Å²) of the DPPC bilayer systems. (E) Mean defect cluster area (Å²) for the two DPPC systems. (F) Mean defect surface coverage, calculated as the total defect divided by the projected membrane area and expressed as a percentage. Error bars in (D-F) represent the standard deviation across three independent replicas.

### 2.3 Mesh Generation

LipiDeA constructs triangulated membrane surfaces independently for each trajectory frame. The membrane surface is defined from a user-selected CG bead type, with GL2 beads used as the default representation of glycerol-based lipid headgroups. Membrane leaflets are first identified using the LeafletFinder algorithm from MDAnalysis based on the mean z-position of beads, which separates upper and lower leaflets^33,34^. For each leaflet, bead coordinates are converted into a point cloud representation. Local surface normals are estimated from neighboring points and oriented consistently such that the upper and lower leaflet normals point in the positive and negative z-direction, respectively. A Poisson surface reconstruction is then applied separately to each leaflet to generate continuous triangular meshes^31,35^.

The resulting meshes are refined through a cleaning procedure that removes duplicated vertices and faces, non-manifold edges, unreferenced vertices, and disconnected components. To ensure numerical consistency in downstream defect calculations, the mesh resolution is further controlled by enforcing a target triangle size distribution defined by the user (mean and standard deviation). This is achieved through iterative mesh refinement and isotropic remeshing using PyMeshLab and Open3D operations^31,36^. Because remeshing modifies connectivity in discrete steps, additional correction cycles are applied when the triangle size distribution deviates beyond the user-defined tolerance. This procedure yields meshes with controlled and approximately uniform triangle areas, which is essential for accurate defect quantification based on triangle-wise area integration.

To avoid artifacts near periodic boundaries, an automatic edge-correction procedure is applied. Candidate boundary triangles are first identified either by convex hull analysis of triangle centroids in the x-y plane or by detecting triangles connected to mesh boundaries. Each candidate triangle is then evaluated for physical support using a KD-tree search to determine whether beads are present near the triangle centroid in the x-y plane^37^. The candidate is retained only if supporting beads are found within the local region and if those beads are consistent with the expected z-position range of the corresponding leaflet. Triangles that lack bead support or whose associated beads fall outside the z-range of both leaflet surfaces are removed. This criterion ensures that only non-physical boundary artifacts are eliminated while preserving physically meaningful edge regions.

Before export, each leaflet mesh undergoes a final quality control step to ensure a topologically consistent surface. Non-manifold vertices and edges introduced during surface reconstruction are automatically corrected. Any small disconnected mesh fragments are then removed by connected component analysis using Open3D, and only the largest connected component is retained for each leaflet^31^. Finally, the reconstructed upper and lower leaflet meshes are converted into numerical representations for downstream analysis. For each frame, LipiDeA stores vertex coordinates, triangle connectivity, triangle normals, and triangle centroids in NPZ format^38^. Mesh files in PLY format are also exported to enable visualization by using molecular graphics software, such as VMD or Chimera^28,29^.

### 2.4 Defect Finding

Lipid packing defects are defined as membrane surface regions that lack sufficient lipid headgroup coverage. For each frame, the centroids of all mesh triangles are first calculated for both membrane leaflets and stored in a KD-tree to enable efficient spatial searches^31,37^. For each lipid bead (target bead), the nearest mesh triangle and its neighboring triangles are identified using a 3D nearest-neighbor search algorithm^31,37^. By default, 48 neighboring triangles are considered, corresponding to a mesh resolution of 1 Å². Candidate lipid beads associated with these local triangles are then collected. To exclude contributions from distant regions of the membrane, candidate beads are retained only when their z-coordinates fall within ±30 Å of the surface along the triangle’s normal. The van der Waals radii of the target bead and all retained candidate beads are then projected onto the local mesh surface.

A barycentric grid is used to determine the fraction of each triangle covered by the projected beads. Phospholipid headgroup coverage is evaluated first: a triangle is classified as a non-defect when any headgroup projection overlaps the triangle; otherwise, it is classified as a defect. Each defect triangle is then assigned to either the neutral-lipid or phospholipid-tail category according to which type covers the larger fraction of the triangle. If the two types of lipids provide equal coverage, the bead with the largest z-coordinate along the triangle normal determines the classification. The resulting defect triangles are exported as a PLY file with different colors for each defect type, so users can visualize and inspect.

### 2.5 Clustering and Deduplication

Defect triangles are grouped into clusters based on mesh connectivity. For each leaflet, defect triangles that share a common edge are assigned to the same defect cluster. Because periodic boundary conditions can produce duplicate representations of the same defect near the simulation box edges, an additional deduplication step is applied. Defect clusters located at periodic boundaries are first wrapped across the simulation box (**Fig. 1(5)**). The distances between cluster centers are then evaluated, and clusters separated by less than 10 Å are considered duplicates. The area of each defect cluster is calculated as the sum of the areas of all triangles belonging to that cluster. Clusters with areas below a user-defined threshold are excluded from further analysis. This filtering step removes small and isolated defects that are unlikely to represent physically meaningful packing defect regions.

### 2.6 Graph-Based Pruning

Some defect clusters remain connected through narrow bridge-like regions composed of only a few triangles (**Fig. 1(6)**). These connections can artificially merge physically distinct packing defects into a single large cluster. To address this issue, LipiDeA applies a graph-based pruning procedure to separate weakly connected defect regions^39^. For each defect cluster, bridge connections are identified using the Tarjan’s bridge-finding algorithm^39^. This algorithm determines whether a connection provides the only path between two regions of a cluster. Connections that do not possess an alternative path are classified as bridges and removed, which separates off the cluster into independent defect regions. Following pruning, cluster areas are recomputed, and the area filter is applied again to remove clusters smaller than the user-defined threshold. The final defect information, e.g., cluster number, cluster area, and 3D mesh data, are reported for visualization and later quantitative defect analysis.

### 2.7 Analysis

LipiDeA provides multiple quantitative analyses for characterizing membrane packing defects. These are organized into three categories: packing-defect statistics computed directly from the defect clusters, monolayer/bilayer classification of each cluster, and curvature analysis of the leaflet surfaces. Users can perform each analysis independently according to their need.

#### 2.7.1 Packing Defect Statistics

The packing defect constant (π) is obtained by fitting the probability distribution of defect areas to an exponential decay model^5,19,40^. First, defect areas from all trajectory frames are grouped into histogram bins to calculate the probability of observing a defect with area *A*. The resulting probability distribution is then fitted to an exponential function,

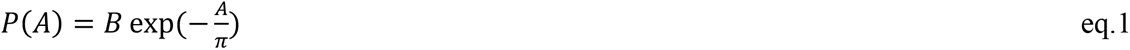

where *B* is the pre-factor, *A* is the defect area, *P(A)* is the probability of the defect with area *A*, and *π* is the packing defect constant. Taking the natural logarithm gives a linear relation,

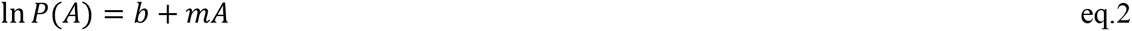

where *b* = ln *B* is the intercept and the *m* is the fitted slope. For an exponential distribution, the packing defect constant is calculated from the fitted slope *π* = −1/*m*. Users can specify the minimum defect area, maximum defect area, and minimum probability included in the fitting procedure. LipiDeA reports the packing defect constant together with the fitted slope, intercept, coefficient of determination (*R*^2^), root-mean-square error (RMSE), and the number of histogram bins used in the fit. The RMSE is calculated as

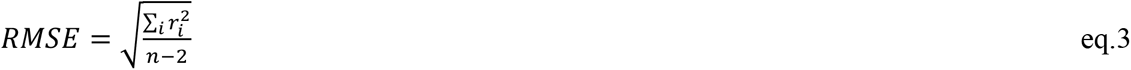

where *n* is the number of fitted histogram bins, and *r_i_* is the residual for bin *i*. The denominator (*n - 2*) accounts for the two fitted parameters, namely the slope and intercept. The residual defines as

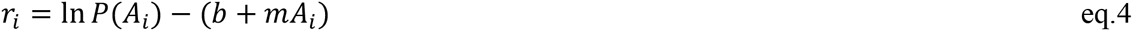

where the first term corresponds to the logarithm of the histogram probability and the second term is the fitted log-probability. The fitting curve and residual statistics are reported to facilitate visual inspection of fit quality. Based on benchmark calculations, a minimum probability cutoff of 10^-^⁴ and trajectories containing at least 2000 frames are recommended to obtain reliable estimates of the packing defect constant.

In addition to the packing defect constant, LipiDeA reports the number of defect clusters and the membrane defect coverage for each trajectory frame. Cluster counts are calculated separately for the upper and lower leaflets and their combined total. Defect coverage is calculated as the total defect area divided by the membrane surface area and is reported for each leaflet, together with the trajectory average. For each frame, the defect area can be further partitioned by defect type, and the neutral lipid and phospholipid tail contributions are reported as fractions of the total defect area. LipiDeA also characterizes the defect size distribution by pooling defect areas from all frames into a histogram and reports the mean defect cluster area.

#### 2.7.2 Monolayer and Bilayer Classification

Membranes containing neutral lipids can contain both bilayer-like and monolayer-like regions within the same system. To distinguish packing defects from these different membrane environments, LipiDeA classifies each defect cluster based on its spatial relationship with the opposite membrane leaflet. For each cluster, the program evaluates the distance between the cluster triangles and the corresponding regions on the opposite leaflet. Users define two parameters for bilayer identification: the maximum distance allowed between the two leaflets (default as 40 Å) and the minimum fraction of cluster triangles that must satisfy this distance criterion (default as 50%). A defect cluster is classified as a bilayer region when the fraction of triangles within the defined leaflet separation distance exceeds the user-defined threshold; otherwise, the cluster is classified as a monolayer region. LipiDeA reports the region classification, cluster area, and paired fraction for each defect cluster. Mesh files separated by monolayer and bilayer regions are also generated for visualization. Optionally, the packing defect constant can be calculated separately for monolayer and bilayer regions to compare their defect properties.

#### 2.7.3 Curvature Analysis

LipiDeA provides membrane curvature analysis to characterize defect properties across different membrane geometries. Local membrane curvature is calculated from the triangulated mesh using the scale-dependent quadric fitting method implemented in PyMeshLab^36^. At each mesh vertex, a local quadric surface is fitted, and the two principal curvatures, *κ*_1_ and *κ*_2_, are obtained. The mean curvature is then calculated as

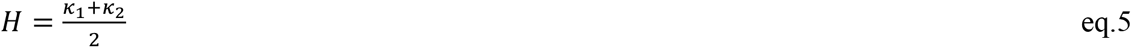

The calculated curvature values are mapped onto the defect clusters identified by LipiDeA. For each defect cluster, the program reports the corresponding curvature values together with cluster area and other defect properties.

## 3. Simulation Settings

### 3.1 System Setup

Two types of membrane systems were applied in this study: 1) DPPC bilayers used to model conventional biological membranes (**Fig. 2A, 2B**) and 2) trilayer assemblies designed to represent LD-like membrane (**Fig. 3**). The initial structures of DPPC bilayer were obtained from Singhal *et al*^23^. and consisted of 3287 DPPC lipids in both planar and buckle geometries. The LD-like systems contain two phospholipid leaflets with compositions of POPC/DOPE/SAPI at a ratio of 65:27:8. A neutral lipid layer, composed of either diacylglycerol (DAG) or triacylglycerol (TAG), is inserted between the two phospholipid layers, as illustrated in **Fig. 3**. To systematically investigate the effect of neutral lipid concentrations on membrane packing defects, a series of systems with varying phospholipid-to-neutral lipid ratios were constructed. Neutral lipid content is expressed relative to the phospholipid composition. For example, a 50% DAG system has a lipid composition of POPC:DOPE:SAPI:DAG = 65:27:8:50 and contains 390 POPC, 162 DOPE, 48 SAPI, and 300 DAG molecules. Analogous TAG systems were generated by replacing DAG with the same number of TAG molecules. The complete series consisted of phospholipid-only membranes (no neutral lipid) and systems containing 1%, 5%, 25%, 50%, 75%, 100%, 125%, and 150% neutral lipid for both DAG and TAG.

**Figure 3:**
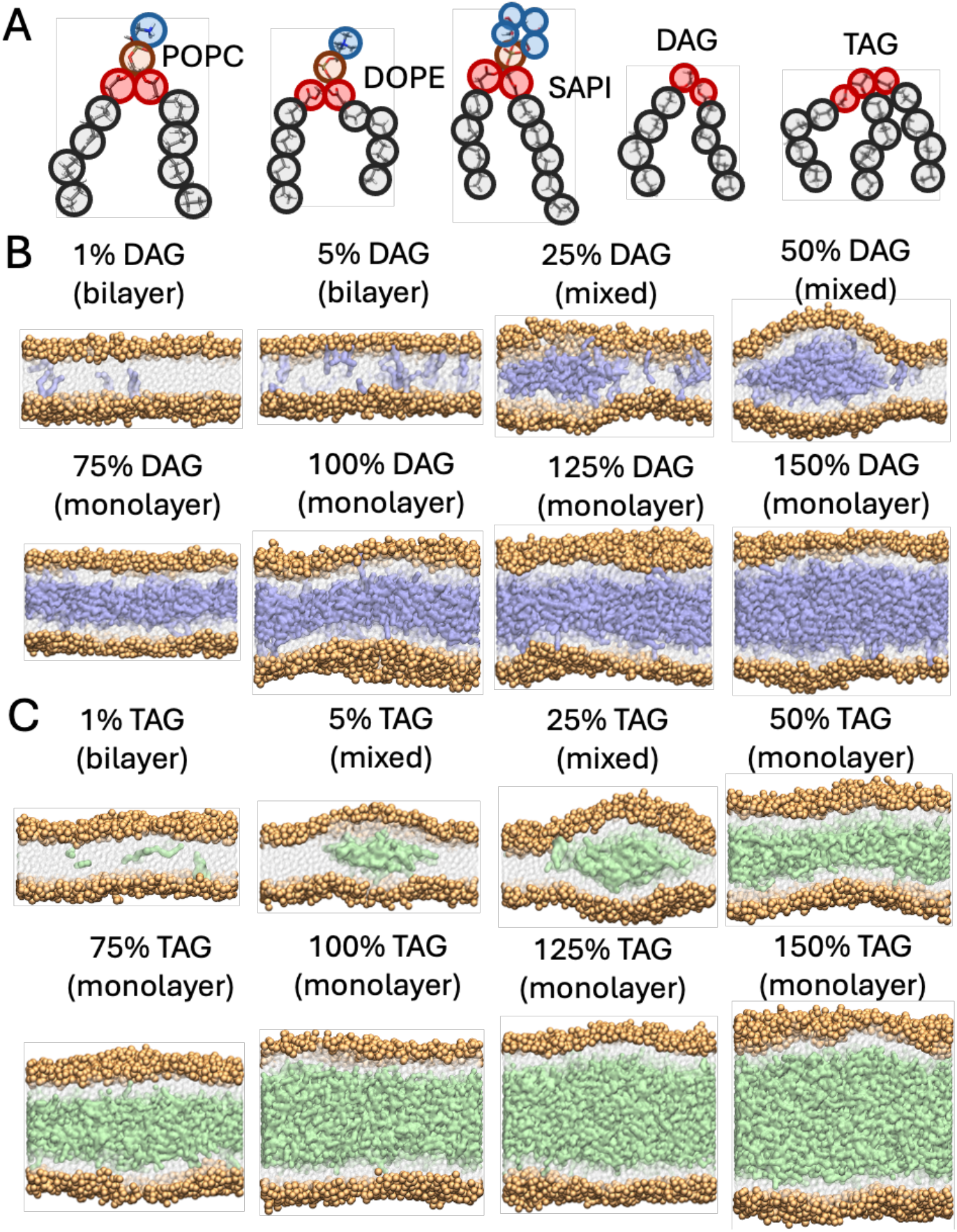
Composition of the LD membrane models. (A) Coarse-grained MARTINI 3 representations of POPC, DOPE, SAPI, DAG, and TAG compared with their corresponding all-atom structures. (B) LD membrane models containing increasing concentration of DAG. (C) LD membrane models containing increasing concentration of TAG. In panels (B) and (C), phospholipid phosphate beads are shown in orange, DAG is blue, TAG is green, and phospholipid tails as white beads.

Initial configurations for all DAG and TAG systems with a neutral lipid ratio <= 50% were generated using CHARMM-GUI^41,42^. For TAG-rich systems with a neutral lipid ratio > 50%, initial coordinates were constructed using PACKMOL to generate a TAG layer, which was then inserted between the two phospholipid leaflets, with an additional 5 nm separation along the z direction between the TAG phase and each phospholipid leaflet to avoid steric overlap prior to equilibration^20,43,44^. Each system was solvated with a 25 Å water layer added above and below the membrane. Sodium and chloride ions were added to achieve a salt concentration of 0.15 M.

### 3.2 Molecular Dynamics Simulations

To compare our results with the work from Singhal *et. al.,* the DPPC bilayer systems were modeled using the MARTINI 2.0 CG force field^23,45^. As the force field of TAG is only available for MARTINI 3.0, we applied it to all LD systems^32,46^. All MD simulations were performed using GROMACS 2018.4^47^. Energy minimization was carried out in two consecutive stages using the steepest descent algorithm, with 5000 steps in each stage^48^. The minimized systems were then equilibrated through six sequential steps with gradually reduced lipid positional restraints and increasing integration time steps. Initial restraints of 200 kJ/mol/nm² were applied with a 2 fs time step. The restraints were subsequently reduced to 100, 50, 20, and 10 kJ/mol/nm², while the corresponding time steps were increased to 5, 10, 15, and 15 fs, respectively. Finally, the restraint was maintained at 10 kJ/mol/nm² for 20 fs. During equilibration, pressure was controlled using the Berendsen barostat in a semi-isotropic ensemble with a coupling constant of 5 ps, a compressibility of 3 × 10^-^⁴ bar^-^¹ in both lateral directions, and a reference pressure of 1 bar^49^.

Production simulations for the DPPC bilayer systems followed the protocol of Singhal *et al*^23^. Briefly, electrostatic and van der Waals interactions were truncated at 1.4 nm, and the temperature was maintained at 325 K. Pressure was controlled using the Parrinello-Rahman barostat with semi-isotropic coupling^50^. For the buckle membrane, the reference pressure was applied only along the z-direction to preserve the imposed membrane curvature. Production simulations for the LD systems were performed at 300 K using the velocity-rescaling thermostat^51^. Pressure was controlled using the Parrinello-Rahman barostat in the semi-isotropic ensemble with a coupling constant of 12 ps, the same compressibility, and the same reference pressure used during equilibration^50^. The cutoff of long-range electrostatic and van der Waals interactions was 1.1 nm. For both the DPPC bilayer and LD systems, simulations were performed under periodic boundary conditions with a relative dielectric constant of 15 and a 20 fs time step. Each system was simulated for 5 μs with three independent replicas.

### 3.3 Post-MD Analysis

The final 2 μs of each trajectory were used for analysis, with frames saved at 1 ns intervals. For mesh generation, the target mean triangle area was set to 1 Å² with a maximum triangle-area coefficient of variation of 0.25. The minimum defect cluster size was set to two triangles. Packing defect constants were calculated from defect area distributions using a linear fit in the 15-250 Å² range. All other LipiDeA parameters were applied using default values. Meshes were inspected using VMD to verify surface reconstruction, and defect distributions and membrane curvature were visualized using VMD and MeshLab^28,30^.

## 4. Results

### 4.1 Validation of LipiDeA Using DPPC Bilayer Systems

To validate LipiDeA, we first analyzed two well-characterized DPPC membrane systems: a planar bilayer and a buckle bilayer (**Fig 2B**). The defect area distributions were fitted to obtain the packing defect constant (**Fig. 2C**). As expected, the buckle membrane exhibited larger packing defects than the planar membrane, with packing defect constants of 21.75 ± 0.29 Å² and 15.61 ± 0.03 Å², respectively (**Fig. 2D**). These values agree well with previous reports, demonstrating that LipiDeA accurately reproduces established packing defect measurements^23^. The buckle membrane also exhibited a larger mean defect area and a greater fraction of membrane surface occupied by packing defects (**Fig. 2E, 2F**). These results are consistent with the larger packing defect constant and indicate greater hydrophobic exposure in the curved buckle membrane.

We next examined how local membrane curvature influences the spatial distribution of packing defects. The planar membrane exhibited a narrow curvature distribution centered near zero, and all defects were located in the regions with mean curvatures within ±0.0025 Å⁻¹ (**Fig. 4A**). In contrast, the buckle membrane sampled a much broader curvature range (**Fig. 4A**), which enabled direct analysis of the relationship between curvature and defect formation. The probability of observing a packing defect increased with positive mean curvature, whereas few defects were detected in negatively curved regions (**Fig. 4B**). Representative surface views further illustrate this trend, with larger and more frequent packing defects observed in positively curved regions compared to flat or negatively curved areas (**Fig. 4C**).

**Figure 4:**
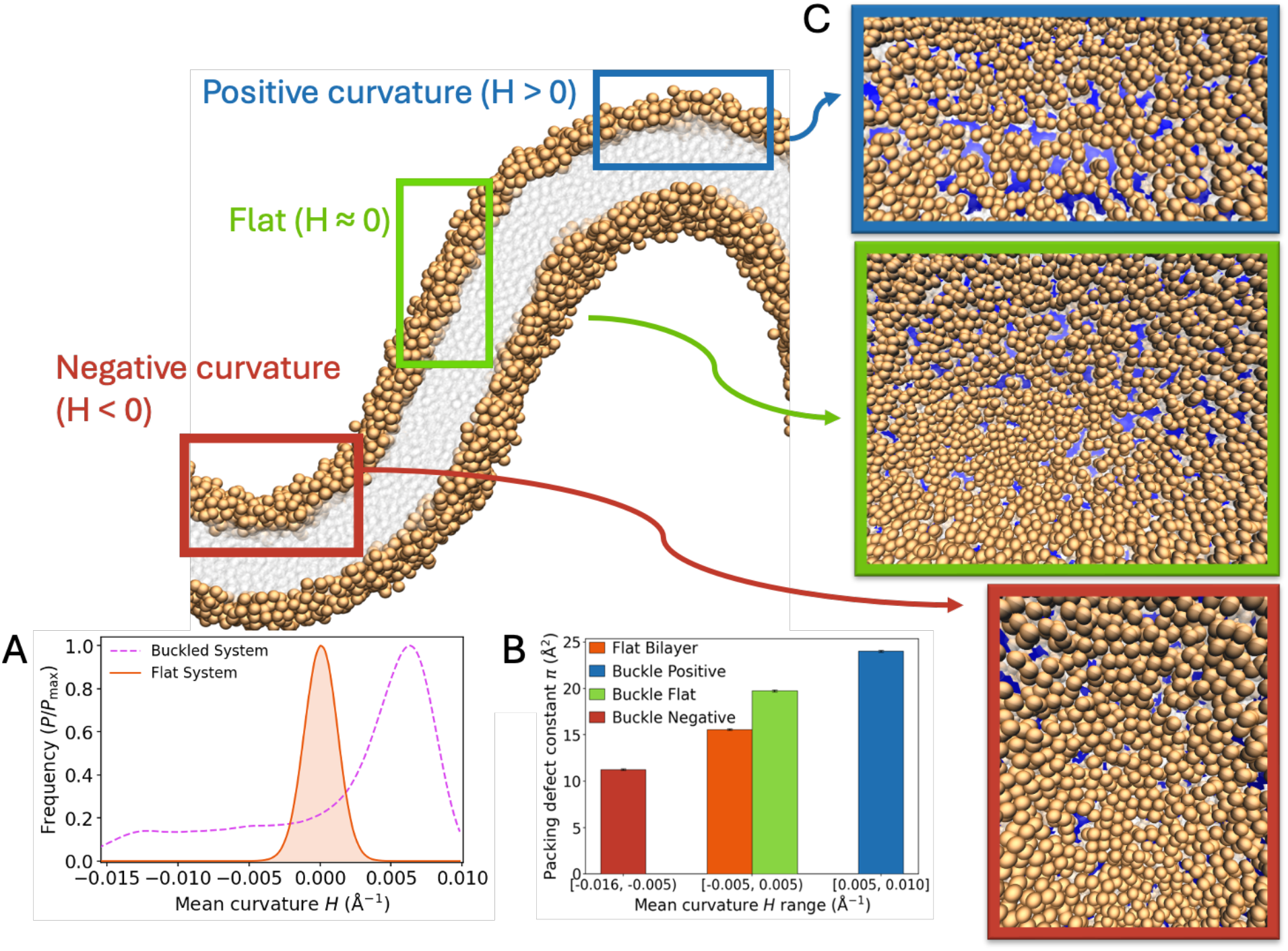
Membrane curvature dependence of Packing defects in DPPC bilayers. (A) Distribution of mean curvature *H (*Å⁻¹) for the planar and buckled membranes. (B) Packing defect constant *π (*Å²), calculated for the three curvatures ranges: negatively curved, near-flat, and positively curved regions. (C) Top views of membrane-surface regions with negative (red), near-zero (green), and positive (blue) curvature.

### 4.2 Application of LipiDeA to Lipid Droplet Membranes

We then applied LipiDeA to LD membrane models containing different concentrations of neutral lipids (**Fig. 3**). Defect area distributions were fitted to obtain the packing defect constant for each system (**Fig. 5A**). Relative to the phospholipid-only membrane (π = 34.41 ± 0.52 Å²), the packing defect constant increased with higher neutral lipid content and reached a maximum at intermediate concentrations (**Fig. 5B**). The largest packing defect constant was observed for the 50% DAG system (52.19 ± 0.06 Å²), whereas the TAG systems reached their maximum at 25% neutral lipids (π = 47.71 ± 0.92 Å²). Beyond these concentrations, the packing defect constant decreased and plateaued as the membranes transitioned into predominantly monolayer structures. Throughout this concentration range, DAG consistently showed larger packing defect constants than TAG. The mean defect area and membrane defect coverage followed the same trend (**Fig. 5C, 5D**), which indicates that intermediate neutral lipid concentrations generate the greatest degree of hydrophobic exposure.

**Figure 5:**
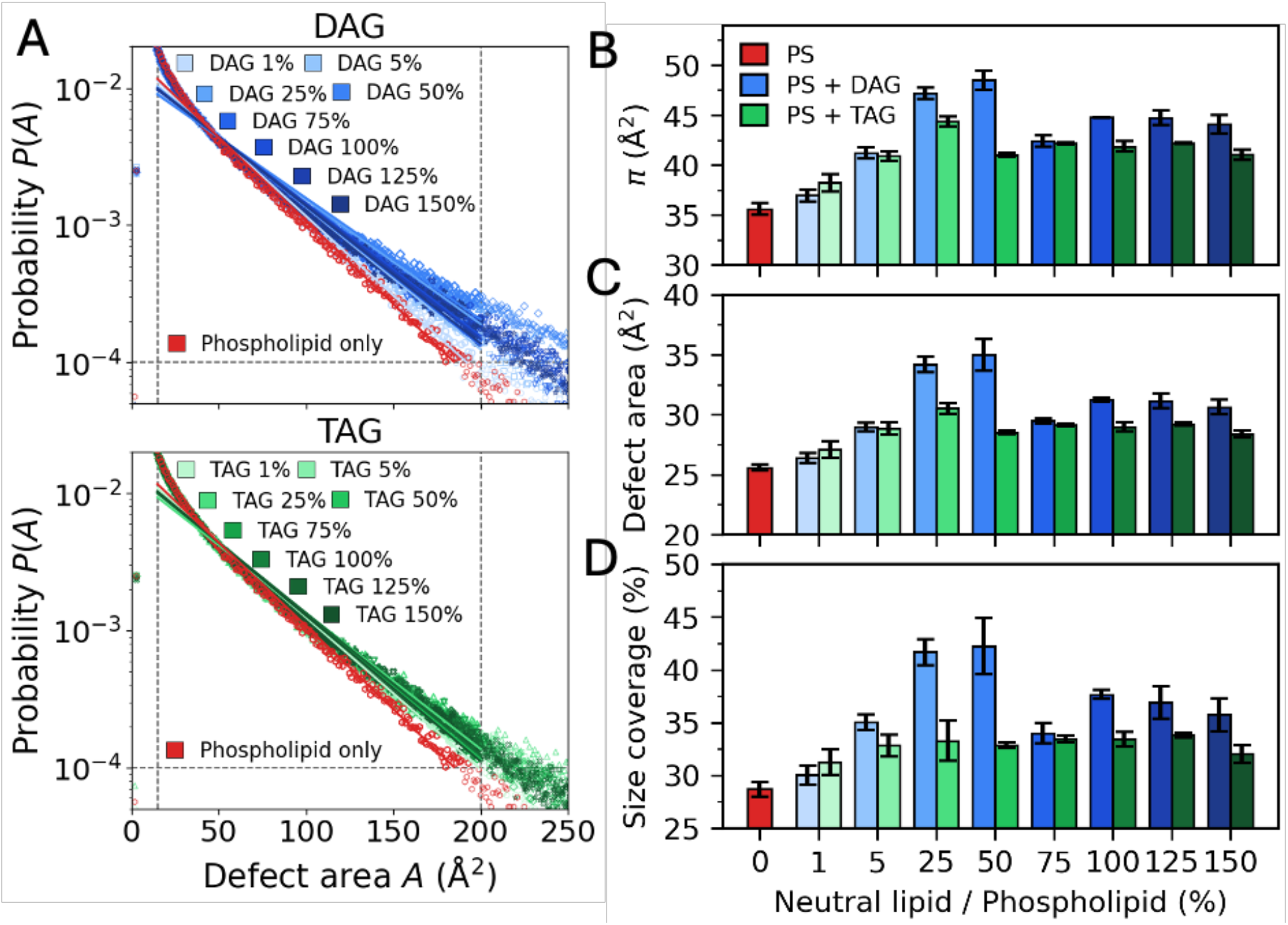
Packing defect statistics for LD membrane models. (A) Defect area probability distributions, *P(A)*, and their exponential fits for the phospholipid-only membrane containing POPC, DOPE, and SAPI (red), DAG-containing membranes (blue), and TAG-containing membranes (green). Fittings were performed over the 15-250 Å² range with a minimum probability cutoff of 10⁻⁴. (B) Packing defect constant, *π (*Å²). (C) mean defect cluster area (Å²). (D) mean defect surface coverage, expressed as the percentage of membrane surface area occupied by defects. Panels (B-D) show the corresponding values for membranes containing different concentrations of neutral lipids.

Because LD systems could contain both bilayer and monolayer regions simultaneously, we used LipiDeA to classify defect clusters according to their local membrane environment. DAG systems formed mixed bilayer-monolayer structures at 25% and 50% neutral lipid concentration, whereas TAG membranes exhibited mixed morphologies at 5% and 25% before becoming complete monolayers at 50% (**Fig. 3**). Within the same membrane, monolayer regions consistently exhibited larger packing defect constants than neighboring bilayer regions (**Fig. 6A**). Once the membranes became complete monolayers at higher neutral lipid concentrations, the packing defect constant decreased, which suggests that the largest packing defects arise during the bilayer-to-monolayer transition rather than in fully developed monolayers.

**Figure 6:**
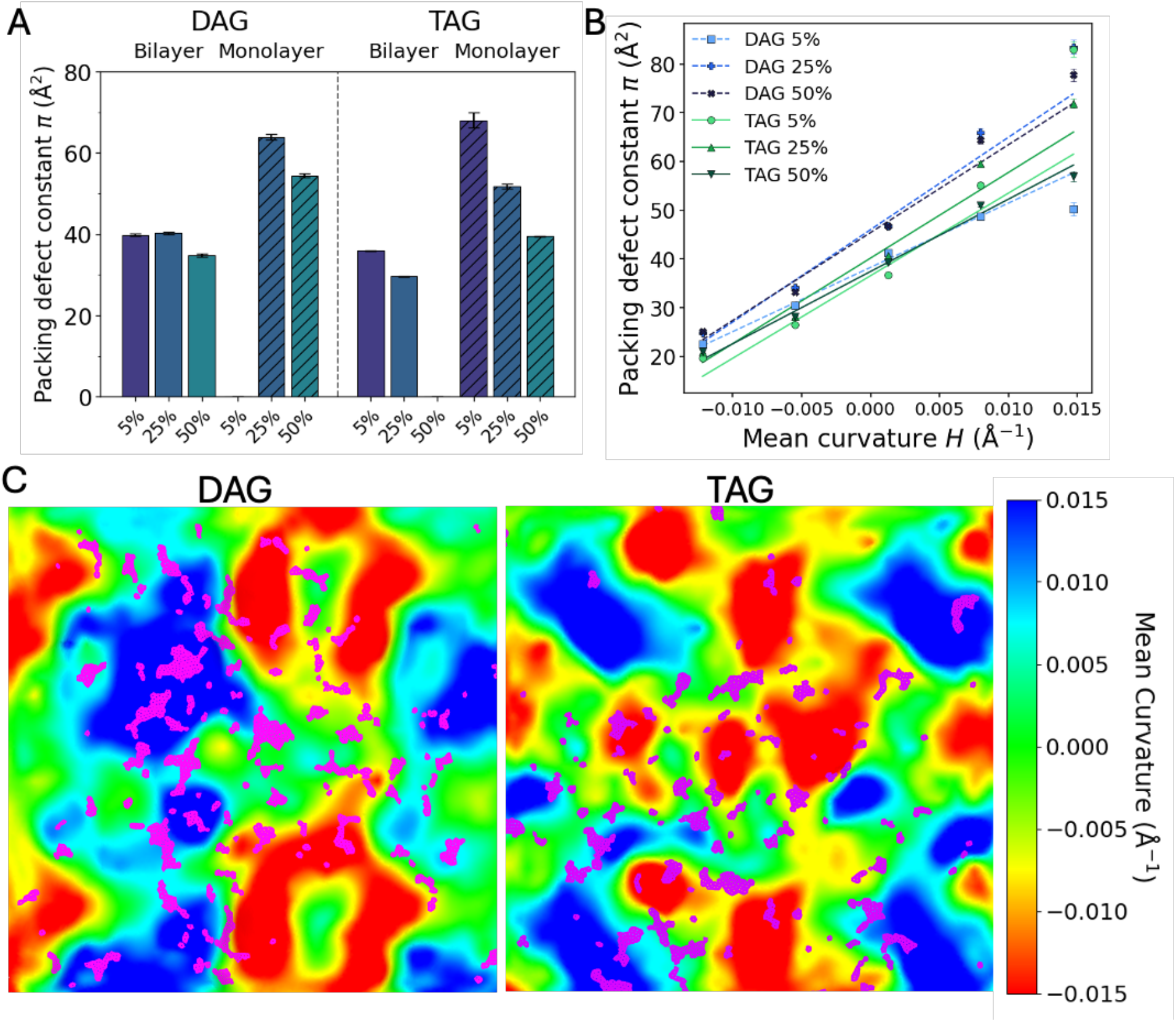
Bilayer/monolayer classification and curvature analysis of the LD membrane models. (A) Packing defect constant, *π (*Å²), calculated separately for defect clusters classified as monolayer or bilayer regions. (B) Packing defect constant calculated across different range of local mean curvature. (C) Curvature heat maps of the membrane surface for the 50% DAG and 50% TAG systems, with identified packing defects shown in magenta. Surface color represents the local mean curvature, *H (*Å⁻¹), ranging from negative (red) through near-flat (green) to positive (blue) curvature.

We next examined how membrane curvature influences packing defect formation in the mixed bilayer-monolayer systems. Across all systems, packing defect size increased with increasing positive mean curvature (**Fig. 6B**), which indicates that positively curved membrane regions are more prone to defect formation. At the same mean curvatures, DAG membranes exhibited larger packing defects than TAG membranes (**Fig. 6C**). However, increasing the neutral lipid concentration beyond the point at which the monolayer formed produced little change in the curvature-defect relationship. These results show that both membrane curvature and lipid composition contribute to packing defect formation, with positive curvature promoting larger defects and DAG membranes exhibiting greater defect propensity than TAG membranes.

LipiDeA also distinguishes packing defects associated with exposed phospholipid tails from those associated with exposed neutral lipids (**Fig. 7A, 7B**). In all systems, most packing defects are contributed from phospholipid tail exposure, which accounted for 90-98% of the total defect area, whereas neutral lipids generally contributed less than 10%. The contribution from neutral lipids increased with increasing neutral lipid concentration, although this increase reached a plateau once the neutral lipid concentration was sufficient to form a lens. DAG-containing systems showed a greater defect contribution from neutral lipids than TAG systems (**Fig. 7B**). DAG also showed a greater tendency to insert between phospholipid headgroups than TAG (**Fig. 7C, 7D**).

**Figure 7:**
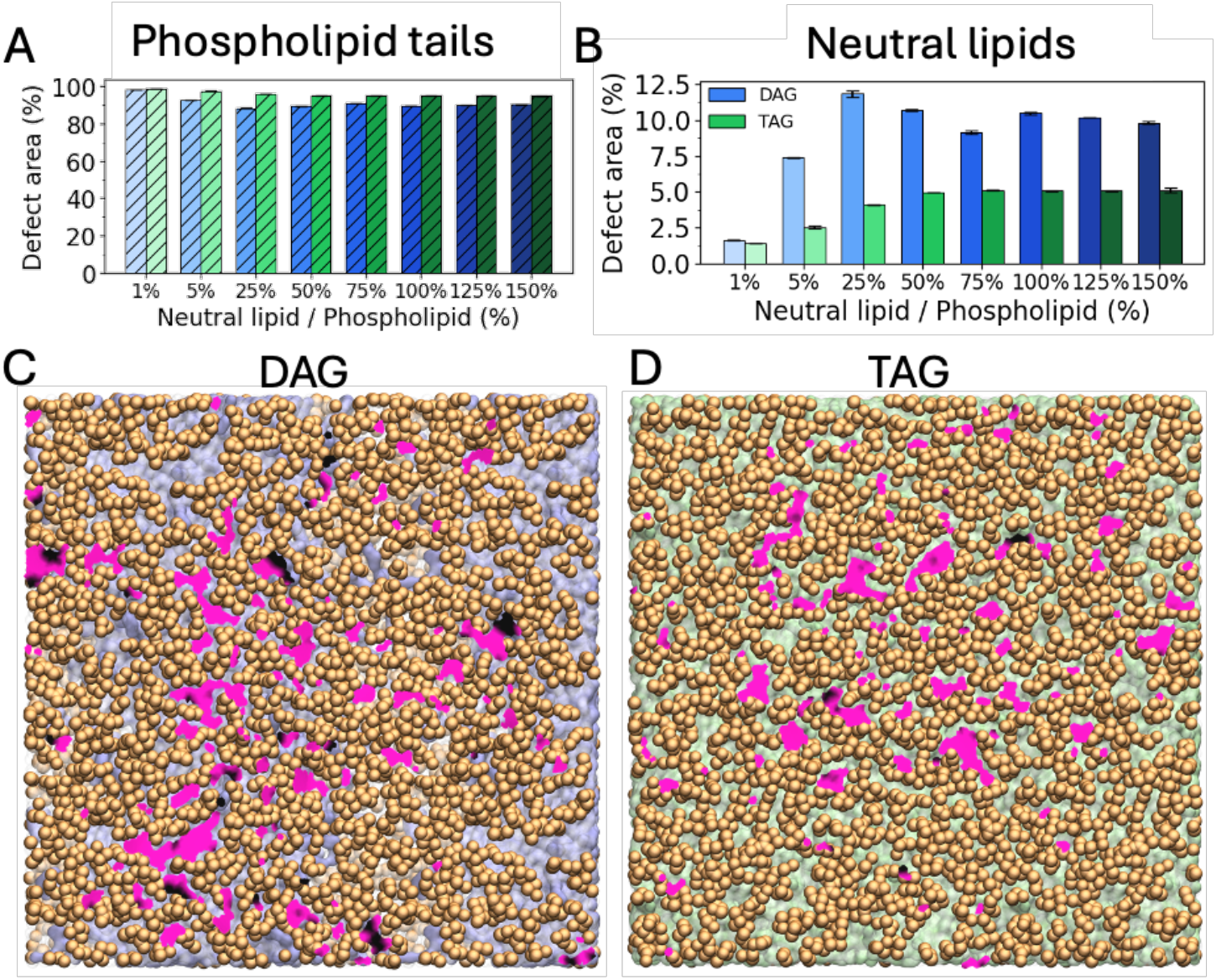
Contribution of phospholipid acyl tails and neutral lipids to packing defects in LD membrane models. (A) Percentage of total defect area contributed by exposed phospholipid acyl tails. (B) Percentage of the total defect area contributed by exposed neutral lipids. (C, D) Top views of the 50% DAG (C) and 50% TAG (D) membrane systems, with defect triangles colored according to defect type: phospholipid acyl tail (magenta) and neutral lipid (black).

### 4.3 Computational Performance

We evaluated the computational performance of LipiDeA across different system sizes and numbers of CPU cores. Two systems were selected for benchmarking: the phospholipid-only system containing 600 lipids of PC/PE/PI and the DPPC buckled system containing 3287 lipids, which represent the smallest and largest systems examined, respectively. All calculations were performed on an AMD Ryzen 7 5800X CPU using 1, 2, 4, or 8 CPU cores. The total computational time and the time required for each LipiDeA step were recorded.

Completion of the full seven-step workflow required 8 hrs for the phospholipid-only system and 90 hrs for the DPPC buckled system when using a single CPU core (**Fig. 8A**). Increasing the number of cores to eight reduced the total computational time to 3 hrs and 36 hrs, respectively. The largest performance improvement occurred when increasing the CPU allocation from one to two cores, whereas additional cores provided progressively smaller reductions in computational time. This observation is consistent with Amdahl’s law, as the overall performance is limited by workflow components that cannot be efficiently parallelized^52^.

**Figure 8.**
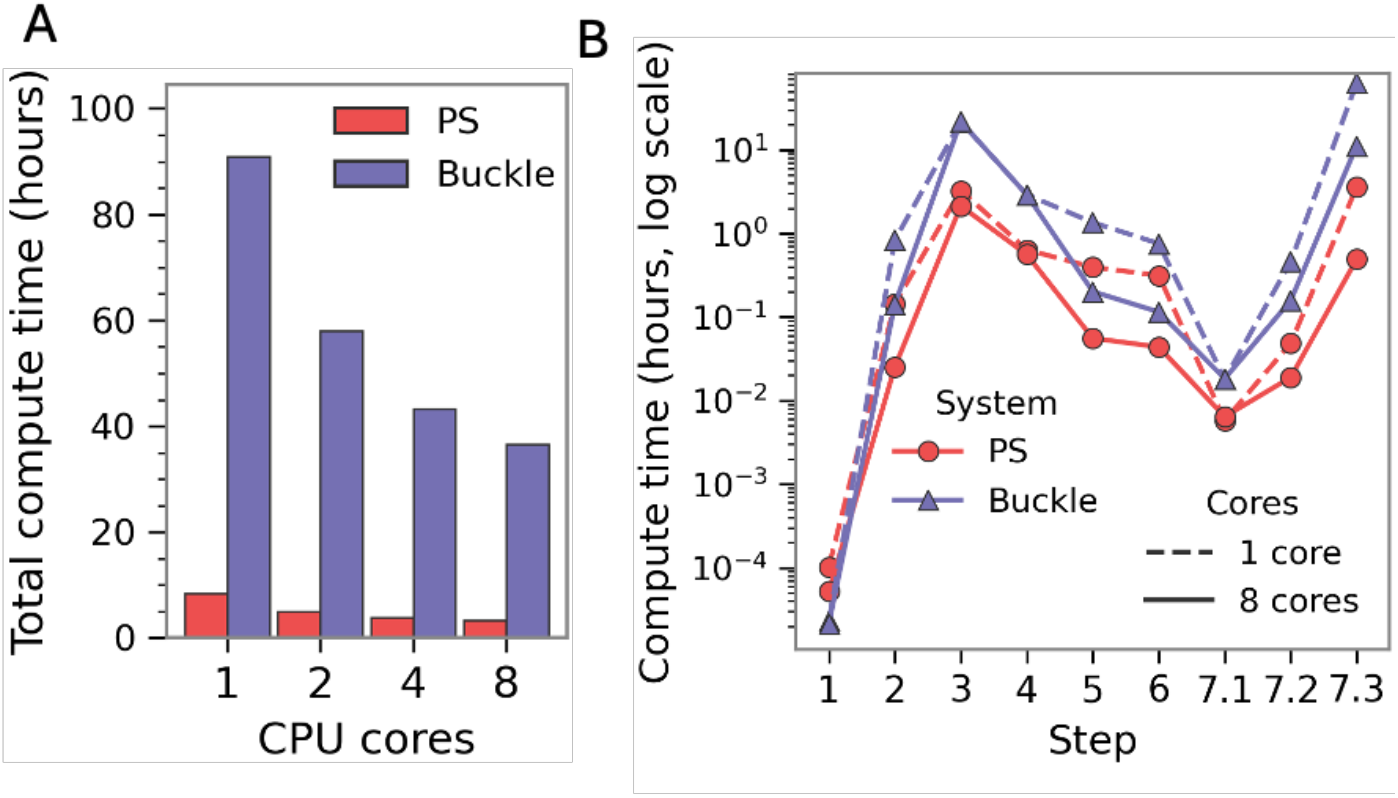
Computational performance of the phospholipid-only and buckled DPPC membrane systems. Computational performance was evaluated for the phospholipid-only membrane containing POPC, DOPE, and SAPI (red) and the buckled DPPC bilayer (purple). (A) Total computational time required to complete Steps 1-7 using different numbers of CPU cores. (B) Computational time required to complete each workflow step using 1 and 8 CPU cores. Computational times are shown on a logarithmic scale.

We then further examined the computational cost of individual LipiDeA steps (**Fig. 8B**). Steps 1 and 7.1 required the least computational time, whereas mesh generation (Step 3) and curvature analysis (Step 7.3) were the most computationally intensive steps. Steps 2, 5, 6, and 7.2 showed improved performance with increasing CPU allocation because these steps use Python-based parallelization and can distribute their workload across multiple CPU cores. In contrast, the computational time for Steps 3 and 4 changed little with the number of cores specified by the user. These steps use internal parallelization provided by Open3D and Numba, respectively, which employ OpenMP-based routines and automatically utilize all available CPU resources^31,53,54^. Consequently, increasing the user-specified cores provides limited additional benefit for these steps.

## 5. Discussion

LipiDeA extends packing defect analysis to heterogeneous membrane systems beyond conventional phospholipid bilayers. Previous methods have successfully quantified packing defects in planar and curved bilayer systems, including grid-based approaches such as PackMem and triangulated-mesh methods such as CG2TS^19,22^. Analysis of the DPPC benchmark systems demonstrated that LipiDeA accurately reproduced previously reported packing defect constants, which supports the validation of both the mesh generation and defect identification algorithms^23^. Beyond reproducing established measurements, the integrated curvature analysis revealed that packing defects preferentially localize to positively curved membrane regions, which provides spatial information that complements the conventional packing defect constant.

Application of LipiDeA to LD membranes demonstrated its ability to characterize heterogeneous membrane environments that contain bilayer-like, monolayer, and neutral-lipid-rich regions within the same system. Increasing neutral lipid content increased the packing defect constant, with the largest defects occurring during the transition from bilayer-dominated to monolayer-containing membranes rather than in fully developed bilayers or monolayers. DAG produced larger packing defects than TAG at comparable concentrations. The smaller and more conical shape of DAG can increase headgroup spacing and disrupt lipid packing at relatively low concentrations, whereas the bulkier TAG molecule remains more compatible with neutral lipid domains until higher concentrations promote lens formation and monolayer development^11,26,27,55,56^. Defect composition analysis further showed that packing defects were dominated by exposed phospholipid tails, whereas neutral-lipid exposure varied with neutral lipid identity and concentration. Curvature analysis also showed that defects preferentially occurred in positively curved regions. These results indicate that packing defects in heterogeneous membranes depend on local membrane organization, composition, and curvature rather than a single global packing defect constant. LipiDeA therefore provides a localized description of membrane packing and hydrophobic exposure in complex membrane systems such as LDs, membrane tubules, and vesicles.

The computational performance analysis highlights both the scalability and practical considerations of LipiDeA. Mesh generation and curvature analysis were the most computationally demanding steps, particularly for large and highly curved membrane systems. The larger computational cost of the buckled DPPC system compared with the phospholipid-only system was primarily associated with its larger and more complex triangulated surface rather than its projected membrane area. This result indicates that computational requirements for LipiDeA depend strongly on mesh complexity, which can increase substantially for corrugated or highly curved membranes. Our modular workflow also provides an important practical advantage for computationally intensive analyses. Because each step depends only on the output of the preceding step, users do not need to regenerate the mesh when modifying downstream analysis parameters, such as defect clustering or fitting ranges. In addition, frame-level and step-level restart capabilities allow interrupted calculations to resume without repeating completed work. These features can reduce the computational cost of analyzing long trajectories.

The mesh triangle size is an important user-defined parameter because it determines both the spatial resolution and computational cost of LipiDeA. Changing the triangle size did not alter the overall trend in packing defects between the planar and buckled DPPC membranes, although larger triangles produced lower packing defect constants (**Fig. 9A**). Smaller triangles provide finer spatial resolution and can capture irregular gaps near lipid headgroups that may be overlooked with larger triangles. In contrast, larger triangles are more readily covered by individual headgroups and can smooth over small defects, although they remain effective for resolving larger defect regions (**Fig. 9C**). Triangle size also strongly affects computational cost, with larger triangles reducing the number of mesh elements and therefore the runtime (**Fig. 9B**). However, excessively large triangles require more trajectory frames to obtain sufficient defect-area samples for reliable packing defect constant estimates. Thus, the triangle size should be selected to balance spatial resolution, computational cost, and the statistical requirements of the analysis.

**Figure 9:**
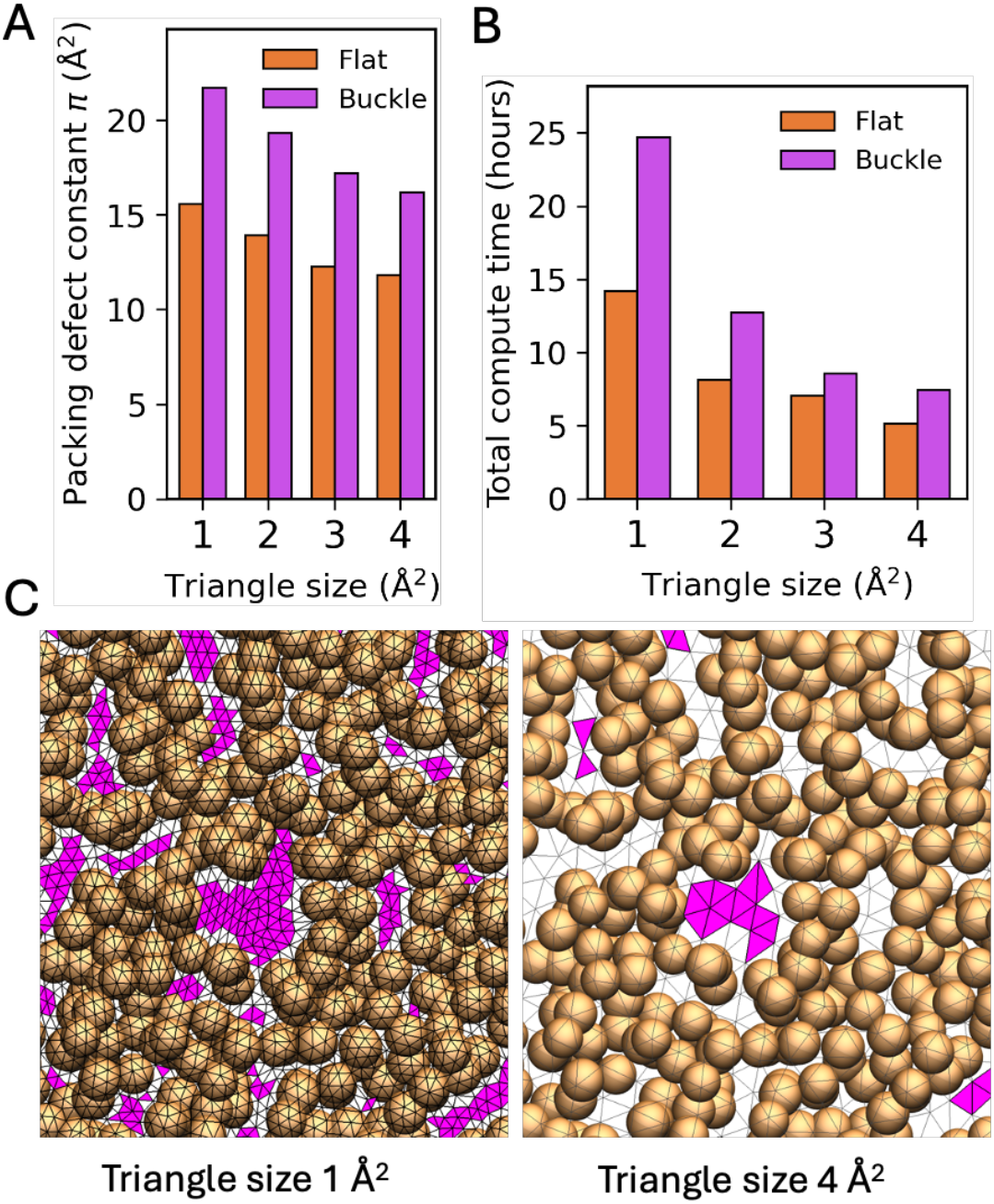
Effect of mesh triangle size on LipiDeA analysis. Results are shown for the planar (orange) and buckled (magenta) DPPC bilayer systems. (A) Packing defect constant, *π (*Å²), calculated using different mesh triangle areas. (B) Computational time required to complete Steps 1-6 with different triangle sizes by using 8 CPU cores. (C) Defect meshes generated with increasing triangle size to show the effect of mesh resolution on resolved defect size and surface coverage.

Several limitations of the current implementation also suggest directions for future development. LipiDeA has been validated primarily for coarse-grained trajectories, and extending the framework to all-atom simulations will require additional testing because of the higher particle density and associated increase in mesh resolution and computational cost. Improving the efficiency of mesh generation and defect identification will therefore be important for applications to large, high-resolution, or highly curved membrane systems. Future development will also extend the current frame-by-frame analysis to track the temporal evolution of individual defect clusters to provide information beyond ensemble-averaged defect statistics. Finally, integrating protein coordinates into the defect analysis would allow packing defects beneath membrane-bound proteins to be characterized directly in terms of their area, composition, curvature, and dynamics. Such an extension could provide a unified framework for relating local membrane packing defects to protein-membrane recognition and binding.

## 6. Conclusion

LipiDeA provides a surface-adaptive approach for quantifying lipid packing defects in membrane systems that extend beyond conventional planar bilayers. It can resolve membrane topology, local curvature, and defect composition enables packing defects to be characterized within heterogeneous membrane environments rather than as a single global property. The application to lipid-droplet-like membranes demonstrates how this approach can reveal changes in membrane packing associated with neutral lipid composition and bilayer-to-monolayer transitions. LipiDeA therefore provides a practical foundation for studying lipid packing defects in increasingly complex membrane structures.

## Data, Materials, and Software Availability

Simulation input files and trajectories have been deposited at https://doi.org/10.5281/zenodo.23020457. The LipiDeA script for membrane packing defects is freely available at https://doi.org/10.5281/zenodo.23020934.

## Acknowledgments

We thank the Wayne State University (WSU) High Performance Computing Center for providing the resources used for CGMD simulations and post-MD analysis. We are grateful to Professor Christopher V. Kelly for helpful discussions and comments, and to Amit Kumar for proofreading the manuscript and offering valuable suggestions on the text. This work was supported by WSU startup funds and NIH grant R35GM160192 (to Y.M.H.).

## TOC

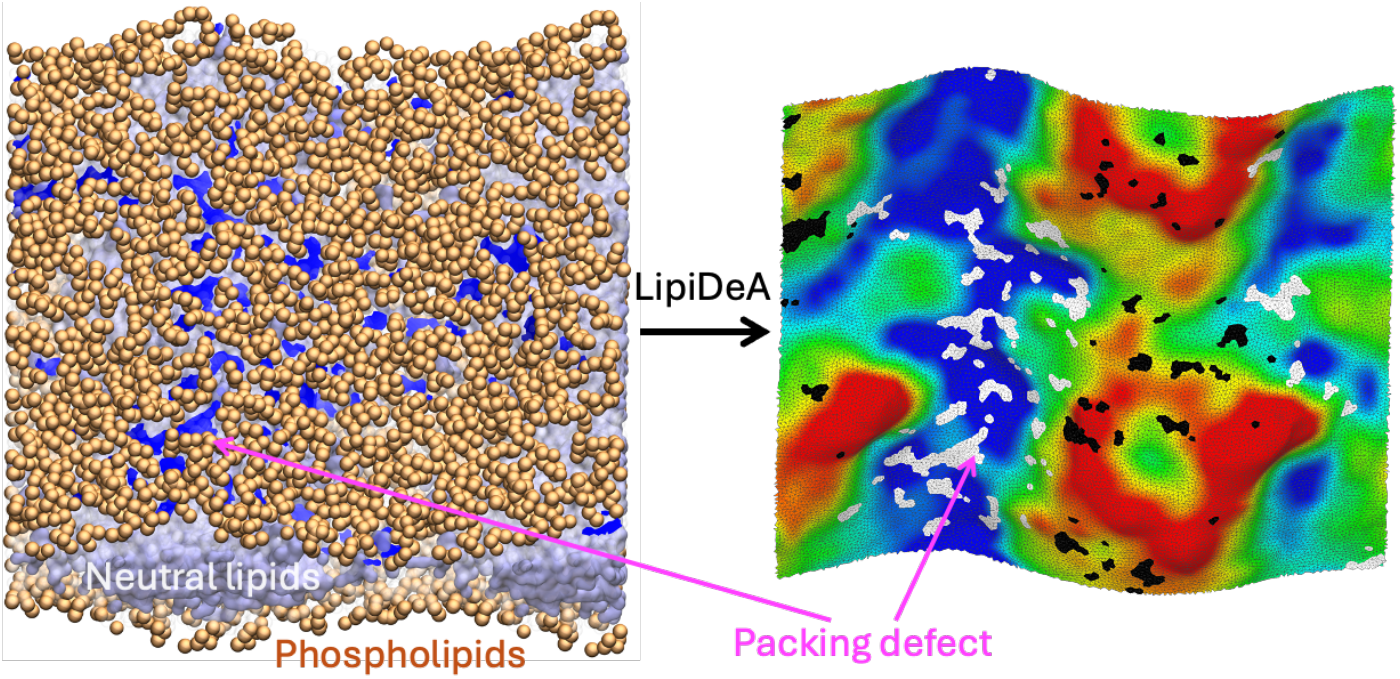

## References

(1) Harayama, T.; Riezman, H. Understanding the Diversity of Membrane Lipid Composition. Nat. Rev. Mol. Cell Biol. 2018, 19 (5), 281–296. 10.1038/nrm.2017.138

(2) Bennett, W. F. D.; Tieleman, D. P. The Importance of Membrane Defects - Lessons from Simulations. Acc. Chem. Res. 2014, 47 (8), 2244–2251. 10.1021/ar4002729

(3) Braun, R. J.; Swanson, J. M. J. Mechanistic Insights into the Selective Targeting of MLX to Triacylglycerol-Rich Lipid Droplets. Biophys. J. 2026, 125 (9), 2084–2098. 10.1016/j.bpj.2026.03.047

(4) Braun, R. J.; Swanson, J. M. J. Progress in Lipid Droplet Simulations: How Polarizability Shapes Triacylglycerol Behavior in Bulk and Interfacial Environments. J. Chem. Theory Comput. 2025, 21 (16), 8235–8248. 10.1021/acs.jctc.5c00882

(5) Vamparys, L.; Gautier, R.; Vanni, S.; Bennett, W. F. D.; Tieleman, D. P.; Antonny, B.; Etchebest, C.; Fuchs, P. F. J. Conical Lipids in Flat Bilayers Induce Packing Defects Similar to That Induced by Positive Curvature. Biophys. J. 2013, 104 (3), 585–593. 10.1016/j.bpj.2012.11.3836

(6) Kim, S.; Oh, M. I.; Swanson, J. M. J. Stressed Lipid Droplets: How Neutral Lipids Relieve Surface Tension and Membrane Expansion Drives Protein Association. J. Phys. Chem. B 2021, 125 (21), 5572–5586. 10.1021/acs.jpcb.1c01795

(7) Chorlay, A.; Thiam, A. R. Neutral Lipids Regulate Amphipathic Helix Affinity for Model Lipid Droplets. J. Cell Biol. 2020, 219 (4), e201907099. 10.1083/jcb.201907099

(8) Turke, M. J.; Raghavan, K. M.; Maltseva, S.; Kerr, D. H. S.; Adams, E. J.; Lee, K. Y. C. Lipid-Packing Defects Are Sufficient to Modulate Membrane Insertion and the Bound State of α-Synuclein. Proc. Natl. Acad. Sci. U.S.A. 2025, 122 (52), e2419823122. 10.1073/pnas.2419823122

(9) Johnson, D. H.; Kou, O. H.; White, J. M.; Ramirez, S. Y.; Margaritakis, A.; Chung, P. J.; Jaeger, V. W.; Zeno, W. F. Lipid Packing Defects Are Necessary and Sufficient for Membrane Binding of α-Synuclein. *Commun*. Biol. 2025, 8 (1), 1179. 10.1038/s42003-025-08622-7

(10) Wildermuth, K. D.; Monje-Galvan, V.; Warburton, L. M.; Klauda, J. B. Effect of Membrane Lipid Packing on Stable Binding of the ALPS Peptide. J. Chem. Theory Comput. 2019, 15 (2), 1418–1429. 10.1021/acs.jctc.8b00945

(11) Tiwari, A.; Pradhan, S.; Sannigrahi, A.; Mahakud, A. K.; Jha, S.; Chattopadhyay, K.; Biswas, M.; Saleem, M. Interplay of Lipid Head Group and Packing Defects in Driving Amyloid-Beta–Mediated Myelin-Like Model Membrane Deformation. J. Biol. Chem. 2023, 299 (5), 104653. 10.1016/j.jbc.2023.104653

(12) Braun, R. J.; Swanson, J. M. J. Capturing the Liquid-Crystalline Phase Transformation: Implications for Protein Targeting to Sterol Ester-Rich Lipid Droplets. Membranes 2022, 12 (10), 949. 10.3390/membranes12100949

(13) Vanni, S.; Hirose, H.; Barelli, H.; Antonny, B.; Gautier, R. A Sub-Nanometre View of How Membrane Curvature and Composition Modulate Lipid Packing and Protein Recruitment. Nat. Commun. 2014, 5 (1), 4916. 10.1038/ncomms5916

(14) Vanni, S.; Vamparys, L.; Gautier, R.; Drin, G.; Etchebest, C.; Fuchs, P. F. J.; Antonny, B. Amphipathic Lipid Packing Sensor Motifs: Probing Bilayer Defects with Hydrophobic Residues. Biophys. J. 2013, 104 (3), 575–584. 10.1016/j.bpj.2012.11.3837

(15) Bhatia, V. K.; Hatzakis, N. S.; Stamou, D. A Unifying Mechanism Accounts for Sensing of Membrane Curvature by BAR Domains, Amphipathic Helices and Membrane-Anchored Proteins. Semin. Cell Dev. Biol. 2010, 21 (4), 381–390. 10.1016/j.semcdb.2009.12.004

(16) Ouberai, M. M.; Wang, J.; Swann, M. J.; Galvagnion, C.; Guilliams, T.; Dobson, C. M.; Welland, M. E. α-Synuclein Senses Lipid Packing Defects and Induces Lateral Expansion of Lipids Leading to Membrane Remodeling. J. Biol. Chem. 2013, 288 (29), 20883–20895. 10.1074/jbc.M113.478297

(17) Campelo, F.; McMahon, H. T.; Kozlov, M. M. The Hydrophobic Insertion Mechanism of Membrane Curvature Generation by Proteins. Biophys. J. 2008, 95 (5), 2325–2339. 10.1529/biophysj.108.133173

(18) Jackson, C. L.; Walch, L.; Verbavatz, J.-M. Lipids and Their Trafficking: An Integral Part of Cellular Organization. Dev. Cell 2016, 39 (2), 139–153. 10.1016/j.devcel.2016.09.030

(19) Gautier, R.; Bacle, A.; Tiberti, M. L.; Fuchs, P. F.; Vanni, S.; Antonny, B. PackMem: A Versatile Tool to Compute and Visualize Interfacial Packing Defects in Lipid Bilayers. Biophys. J. 2018, 115 (3), 436–444. 10.1016/j.bpj.2018.06.025

(20) Kim, S.; Swanson, J. M. J. The Surface and Hydration Properties of Lipid Droplets. Biophys. J. 2020, 119 (10), 1958–1969. 10.1016/j.bpj.2020.10.001

(21) Tripathy, M.; Thangamani, S.; Srivastava, A. Three-Dimensional Packing Defects in Lipid Membrane as a Function of Membrane Order. J. Chem. Theory Comput. 2020, 16 (12), 7800–7816. 10.1021/acs.jctc.0c00609

(22) van der Pol, R. W. I.; Brinkmann, B. W.; Sevink, G. J. A. Analyzing Lipid Membrane Defects via a Coarse-Grained to Triangulated Surface Map: The Role of Lipid Order and Local Curvature in Molecular Binding. J. Chem. Theory Comput. 2024, 20 (7), 2888–2900. 10.1021/acs.jctc.4c00082

(23) Singhal, A.; Van Daalen, T. J. J.; Sevink, G. J. A. The Significance of Lipid Membrane Curvature and Lipid Order for Nano-C60 Uptake. Nanoscale 2025, 17 (41), 24325–24342. 10.1039/D5NR02822A

(24) Kim, S.; Swanson, J. M. J.; Voth, G. A. Computational Studies of Lipid Droplets. J. Phys. Chem. B 2022, 126 (11), 2145–2154. 10.1021/acs.jpcb.2c00292

(25) Sapia, J.; Vanni, S. Molecular Dynamics Simulations of Intracellular Lipid Droplets: A New Tool in the Toolbox. FEBS Lett. 2024, 598 (10), 1143–1153. 10.1002/1873-3468.14879

(26) Campomanes, P.; Zoni, V.; Vanni, S. Local Accumulation of Diacylglycerol Alters Membrane Properties Nonlinearly Due to Its Transbilayer Activity. Commun. Chem. 2019, 2 (1), 72. 10.1038/s42004-019-0175-7

(27) Zoni, V.; Khaddaj, R.; Campomanes, P.; Thiam, A. R.; Schneiter, R.; Vanni, S. Pre-Existing Bilayer Stresses Modulate Triglyceride Accumulation in the ER Versus Lipid Droplets. eLife 2021, 10, e62886. 10.7554/eLife.62886

(28) Humphrey, W.; Dalke, A.; Schulten, K. VMD: Visual Molecular Dynamics. J. Mol. Graphics 1996, 14 (1), 33–38. 10.1016/0263-7855(96)00018-5

(29) Pettersen, E. F.; Goddard, T. D.; Huang, C. C.; Couch, G. S.; Greenblatt, D. M.; Meng, E. C.; Ferrin, T. E. UCSF Chimera—A Visualization System for Exploratory Research and Analysis. J. Comput. Chem. 2004, 25 (13), 1605–1612. 10.1002/jcc.20084

(30) Cignoni, P.; Callieri, M.; Corsini, M.; Dellepiane, M.; Ganovelli, F.; Ranzuglia, G. MeshLab: An Open-Source Mesh Processing Tool. In Eurographics Italian Chapter Conference; Scarano, V., Chiara, R. D., Erra, U., Eds.; The Eurographics Association, 2008. 10.2312/LocalChapterEvents/ItalChap/ItalianChapConf2008/129-136

(31) Zhou, Q.-Y.; Park, J.; Koltun, V. Open3D: A Modern Library for 3D Data Processing. arXiv:1801.09847 2018.

(32) Souza, P. C. T.; Alessandri, R.; Barnoud, J.; Thallmair, S.; Faustino, I.; Grünewald, F.; Patmanidis, I.; Abdizadeh, H.; Bruininks, B. M. H.; Wassenaar, T. A.; Kroon, P. C.; Melcr, J.; Nieto, V.; Corradi, V.; Khan, H. M.; Domański, J.; Javanainen, M.; Martinez-Seara, H.; Reuter, N.; Best, R. B.; Vattulainen, I.; Monticelli, L.; Periole, X.; Tieleman, D. P.; de Vries, A. H.; Marrink, S. J. Martini 3: A General Purpose Force Field for Coarse-Grained Molecular Dynamics. Nat. Methods 2021, 18 (4), 382–388. 10.1038/s41592-021-01098-3

(33) Gowers, R.; Linke, M.; Barnoud, J.; Reddy, T.; Melo, M.; Seyler, S.; Domański, J.; Dotson, D.; Buchoux, S.; Kenney, I.; Beckstein, O. MDAnalysis: A Python Package for the Rapid Analysis of Molecular Dynamics Simulations. In Proceedings of the 15th Python in Science Conference; SciPy: Austin, TX, 2016; pp 98–105. 10.25080/Majora-629e541a-00e

(34) Michaud-Agrawal, N.; Denning, E. J.; Woolf, T. B.; Beckstein, O. MDAnalysis: A Toolkit for the Analysis of Molecular Dynamics Simulations. J. Comput. Chem. 2011, 32 (10), 2319–2327. 10.1002/jcc.21787

(35) Kazhdan, M.; Bolitho, M.; Hoppe, H. Poisson Surface Reconstruction. In Proceedings of the Fourth Eurographics Symposium on Geometry Processing; SGP ’06; Eurographics Association: Aire-la-Ville, Switzerland, 2006; pp 61–70. 10.2312/SGP/SGP06/061-070

(36) Muntoni, A.; Cignoni, P. PyMeshLab, 2021. 10.5281/zenodo.4438750

(37) Virtanen, P.; Gommers, R.; Oliphant, T. E.; Haberland, M.; Reddy, T.; Cournapeau, D.; Burovski, E.; Peterson, P.; Weckesser, W.; Bright, J.; van der Walt, S. J.; Brett, M.; Wilson, J.; Millman, K. J.; Mayorov, N.; Nelson, A. R. J.; Jones, E.; Kern, R.; Larson, E.; Carey, C. J.; Polat, İ.; Feng, Y.; Moore, E. W.; VanderPlas, J.; Laxalde, D.; Perktold, J.; Cimrman, R.; Henriksen, I.; Quintero, E. A.; Harris, C. R.; Archibald, A. M.; Ribeiro, A. H.; Pedregosa, F.; van Mulbregt, P. SciPy 1.0: Fundamental Algorithms for Scientific Computing in Python. Nat. Methods 2020, 17 (3), 261–272. 10.1038/s41592-019-0686-2

(38) Harris, C. R.; Millman, K. J.; van der Walt, S. J.; Gommers, R.; Virtanen, P.; Cournapeau, D.; Wieser, E.; Taylor, J.; Berg, S.; Smith, N. J.; Kern, R.; Picus, M.; Hoyer, S.; van Kerkwijk, M. H.; Brett, M.; Haldane, A.; del Río, J. F.; Wiebe, M.; Peterson, P.; Gérard-Marchant, P.; Sheppard, K.; Reddy, T.; Weckesser, W.; Abbasi, H.; Gohlke, C.; Oliphant, T. E. Array Programming with NumPy. Nature 2020, 585 (7825), 357–362. 10.1038/s41586-020-2649-2

(39) Tarjan, R. E. A Note on Finding the Bridges of a Graph. Inf. Process. Lett. 1974, 2 (6), 160–161. 10.1016/0020-0190(74)90003-9

(40) Cui, H.; Lyman, E.; Voth, G. A. Mechanism of Membrane Curvature Sensing by Amphipathic Helix Containing Proteins. Biophys. J. 2011, 100 (5), 1271–1279. 10.1016/j.bpj.2011.01.036

(41) Qi, Y.; Ingólfsson, H. I.; Cheng, X.; Lee, J.; Marrink, S. J.; Im, W. CHARMM-GUI Martini Maker for Coarse-Grained Simulations with the Martini Force Field. J. Chem. Theory Comput. 2015, 11 (9), 4486–4494. 10.1021/acs.jctc.5b00513

(42) Wu, E. L.; Cheng, X.; Jo, S.; Rui, H.; Song, K. C.; Dávila-Contreras, E. M.; Qi, Y.; Lee, J.; Monje-Galvan, V.; Venable, R. M.; Klauda, J. B.; Im, W. CHARMM-GUI Membrane Builder Toward Realistic Biological Membrane Simulations. J. Comput. Chem. 2014, 35 (27), 1997–2004. 10.1002/jcc.23702

(43) Kumar, A.; Sanders, M.; Zhang, H.; Zhou, L.; Parveen, S.; Jensen, M. C.; Jørgensen, T. J. D.; Kelly, C. V.; Granneman, J. G.; Huang, Y. M. Unraveling the Molecular Mechanisms of ABHD5 Membrane Targeting. eLife 2026, 15, RP109517. 10.7554/eLife.109517

(44) Martínez, L.; Andrade, R.; Birgin, E. G.; Martínez, J. M. PACKMOL: A Package for Building Initial Configurations for Molecular Dynamics Simulations. J. Comput. Chem. 2009, 30 (13), 2157–2164. 10.1002/jcc.21224

(45) Marrink, S. J.; Risselada, H. J.; Yefimov, S.; Tieleman, D. P.; De Vries, A. H. The MARTINI Force Field: Coarse Grained Model for Biomolecular Simulations. J. Phys. Chem. B 2007, 111 (27), 7812–7824. 10.1021/jp071097f

(46) Biasi, L. C. K.; Rodrigues, C. E. da C.; Pessoa Filho, P. de A. Molecular Dynamics of Vegetable Oil Extraction and Degumming: Analysis of Micelles and Phospholipid Bilayers in Different Solvents. Journal of the American Oil Chemists’ Society 2024, 101 (2), 205–223. 10.1002/aocs.12731

(47) Van Der Spoel, D.; Lindahl, E.; Hess, B.; Groenhof, G.; Mark, A. E.; Berendsen, H. J. C. GROMACS: Fast, Flexible, and Free. J. Comput. Chem. 2005, 26 (16), 1701–1718. 10.1002/jcc.20291

(48) Meza, J. C. Steepest Descent. WIREs Comput. Stat. 2010, 2 (6), 719–722. 10.1002/wics.117

(49) Berendsen, H. J. C.; Postma, J. P. M.; van Gunsteren, W. F.; DiNola, A.; Haak, J. R. Molecular Dynamics with Coupling to an External Bath. J. Chem. Phys. 1984, 81 (8), 3684–3690. 10.1063/1.448118

(50) Parrinello, M.; Rahman, A. Polymorphic Transitions in Single Crystals: A New Molecular Dynamics Method. J. Appl. Phys. 1981, 52 (12), 7182–7190. 10.1063/1.328693

(51) Bussi, G.; Donadio, D.; Parrinello, M. Canonical Sampling Through Velocity Rescaling. J. Chem. Phys. 2007, 126 (1), 014101. 10.1063/1.2408420

(52) Amdahl, G. M. Validity of the Single Processor Approach to Achieving Large Scale Computing Capabilities. In Proceedings of the April 18–20, 1967, Spring Joint Computer Conference; ACM: New York, NY, 1967; pp 483–485. 10.1145/1465482.1465560

(53) Dagum, L.; Menon, R. OpenMP: An Industry Standard API for Shared-Memory Programming. IEEE Comput. Sci. Eng. 1998, 5 (1), 46–55. 10.1109/99.660313

(54) Lam, S. K.; Pitrou, A.; Seibert, S. Numba: A LLVM-Based Python JIT Compiler. In Proceedings of the Second Workshop on the LLVM Compiler Infrastructure in HPC; ACM: New York, NY, 2015; pp 1–6. 10.1145/2833157.2833162

(55) Alwarawrah, M.; Dai, J.; Huang, J. Modification of Lipid Bilayer Structure by Diacylglycerol: A Comparative Study of Diacylglycerol and Cholesterol. J. Chem. Theory Comput. 2012, 8 (2), 749–758. 10.1021/ct200790q

(56) Alwarawrah, M.; Hussain, F.; Huang, J. Alteration of Lipid Membrane Structure and Dynamics by Diacylglycerols with Unsaturated Chains. Biochim. Biophys. Acta, Biomembr. 2016, 1858 (2), 253–263. 10.1016/j.bbamem.2015.11.014

